# Modulation of autism-associated serotonin transporters by palmitoylation: Insights into the molecular pathogenesis and targeted therapies for autism spectrum disorder

**DOI:** 10.1101/2025.03.12.642908

**Authors:** Christopher R. Brown, James D. Foster

**Affiliations:** Department of Biomedical Sciences, University of North Dakota School of Medicine and Health Sciences, Grand Forks, ND 58202-9037

**Keywords:** Serotonin Transporter, Autism Spectrum Disorder, Selective Serotonin Reuptake Inhibitor, Post Translational Modification, Protein Palmitoylation

## Abstract

**Background:** Autism spectrum disorder (ASD) is a developmental disorder of the nervous system characterized by a deficiency in interpersonal communication skills, a pathologic tendency for repetitive behaviors, and highly restrictive interests. The spectrum is a gradient-based construct used to categorize the widely varying degrees of ASD phenotypes, and has been linked to a genetic etiology in 25% of cases. Prior studies have revealed that 30% of ASD patients exhibit hyperserotonemia, or elevated whole blood serotonin, implicating the serotonergic system in the pathogenesis of ASD. Likewise, escitalopram, a selective-serotonin reuptake inhibitor (SSRI), has been demonstrated to improve aberrant behavior and irritability in ASD patients, potentially by modulating abnormal brain activation. Prior studies have uncovered proband patients with rare mutations in the human serotonin transporter (hSERT) that manifest enhanced surface expression and transport capacity, suggesting that abnormal enhancement of hSERT function may be involved in the pathogenesis of ASD.

**Methods:** HEK-293 cells stably expressing WT, C109A, I425L, F465L, L550V, or K605N hSERT were subject to analysis for palmitoylation via Acyl-Biotin Exchange followed with hSERT immunoblotting. F465L functional enhancement was confirmed by surface analysis via biotinylation and saturation analysis via 5HT transport. F465L palmitoylation, surface expression and transport capacity were then assessed following treatment with 2-bromopalmitate or escitalopram.

**Results:** Here, we reveal that palmitoylation is enhanced in the ASD hSERT F465L and L550V coding variants, and confirm prior reports of enhanced kinetic activity and surface expression of F465L. Subsequently, treatment of F465L with the irreversible palmitoyl acyl-transferase inhibitor, 2-bromopalmitate (2BP), or escitalopram, rectified enhanced F465L palmitoylation, surface expression, and transport capacity to basal WT levels.

**Limitations:** Tests assessing L550V for surface expression, transport capacity, and reactivity to inhibition of palmitoylation was not assessed. In addition, further characterization is necessary for internalization rates, degradative mechanisms, the impact of cysteine-mediated substitutions, and other SSRIs on these processes.

**Conclusions:** Overall, our results implicate disordered hSERT palmitoylation in the pathogenesis of serotonergic ASD subtypes, with basal recovery of these processes following escitalopram providing insight into its molecular utility as an ASD therapeutic.

## BACKGROUND

Autism spectrum disorder (ASD) is a developmental disorder of the nervous system characterized by a deficiency in interpersonal communication skills, a pathologic tendency for repetitive behaviors, and highly restrictive interests (1). ASD is an increasingly diagnosed condition, demonstrating prevalence across the globe and all socio-economic strata (2). The United States (US) has an increased prevalence of 0.92% when compared to other countries, with Caucasian children being the most commonly affected (1, 3, 4). In the US, one out of every 59 children are diagnosed with ASD, with males outpacing females at a ratio of 4:1 (1, 3, 5–7). The high prevalence of this disorder is acknowledged, in part, to the ‘autism spectrum’ which includes all individuals that exemplify core phenotypic traits. This spectrum includes severe features and more mild forms like those of the previous DSM-4 diagnosis of Asperger’s syndrome (8). The diverse epidemiology (9, 10), histological (10–12), and symptomatic (13) findings of ASD suggest that it’s pathogenesis is complex and multifaceted. Indeed, it is widely accepted that ASD is manifested by a variety of genetic and environmental factors that broadly impact neurobiological development (14–16).

In the early years of ASD research, investigations for a biomarker revealed that 30% of patients developed hyperserotonemia, or elevated whole blood serotonin (5-hydroxytryptamine [5HT]) (17–19). This biomarker is still in use today at some centers as an adjunct in ASD diagnosis (20, 21). In the central nervous system (CNS), 5HT is a monoamine neurotransmitter that modulates sleep, mood, and cognition. Following its release from the pre-synaptic neuron, the extra-neuronal concentration of 5HT is tightly controlled in a spatial and temporal manner by the human serotonin transporter (hSERT). Outside of the CNS, hSERT is expressed in the gastrointestinal tract (22), placenta (23), lung (24), lymphocytes (25), and thrombocytes (blood platelets) (26–29). Originating from enterochromaffin cells of the gut, peripheral 5HT is released into the circulation and sequestered by thrombocytes *via* hSERT, and is reflective of nearly all whole-blood 5HT (30). As with other serotonergic tissue, hSERT is expressed from a single gene (*Locus 17q11.2-21; SLCA4 gene*) in the megakaryocyte, the thrombocyte precursor, before physiologic pyknosis and cellular maturation (31, 32). Importantly, this suggests that defects in hSERT or other 5HT machinery may be involved in the pathogenesis of ASD and is reflected by alterations in whole-blood 5HT levels (30, 33).

Subsequent investigations for genetic links in ASD proband families revealed several coding variants in exons for hSERT. Sutcliffe and colleagues identified five hSERT coding variants: Gly56Ala (G56A), Ile425L (I425L), Phe465Leu (F465L), Leu550Val (L550V), and Lys605Asn (K605N) (34). With the exception of K605N, each coding variant has been associated with typical ASD symptomology including obsessive tendencies, compulsive behavior, resistance to trivial change, and a pathologic attachment to immaterial objects (35, 36). Functional analyses of these variants have revealed an enhancement in 5HT transport capacity and an elevation in cell surface expression for I425L, F465L, and L550V (30, 37), suggesting that these mutations confer a reduction in serotonergic tone. Functional enhancement of the N- and C-termini coding variants, G56A and K605N, have been demonstrated to lack sensitivity to modulation of cyclic GMP-dependent protein kinase (PKG) and p38 mitogen-activated protein kinase (p38 MAPK) pathways (37–39). It has been inferred that SERT surface expression is driven when these pathways are stimulated, suggesting that enhanced activity of these variants results from direct catalytic enhancement by the mutation or disruptions in kinase and/or phosphatase recognition of SERT (37). In comparison, coding variants with mutations in transmembrane domain regions of SERT (I425L, F465L, and L550V) confer an elevation in SERT surface expression and transport capacity, suggesting that mutational alteration of SERT’s structure increases recruitment to or retention at the cell surface through an unknown mechanism (30). Clinically, It has been demonstrated that the rigid-compulsive trait domain is influenced strongly by these enhanced variants (35), which is consistent with clinical evidence that inhibition by selective-serotonin reuptake inhibitors (SSRIs) provide relief for anxiety-like features in adults with ASD (40, 41). Studies have found that abnormal brain activation patterns of patients with ASD (42–44) was abolished following citalopram and/or fluoxetine dosing in patients (43, 44). Furthermore, escitalopram was found in one study to improve aberrant behavior and irritability in ASD patients from baseline to the six-week endpoint (45). These results suggest that many SSRI subtypes may improve ASD symptomatology, possibly by modulating abnormal brain activation.

hSERT function is regulated by numerous signaling pathways that serve to maintain 5HT homeostasis in accordance with physiological demands. Evidence suggests that hSERT is modified *via* post-translational modifications (PTM) like phosphorylation (46–48), glycosylation (49, 50), and ubiquitylation (51, 52) that serve to coordinate SERT kinetic activity, localization, and surface expression in response to cellular requirements. Previously, SERT has been identified as a target for the lipid-based post-translational modification, S-palmitoylation (53–55). Analogous to phosphorylation, palmitoylation is dynamic and reversible, controlling protein folding, activity, trafficking and localization in accordance with physiologic requirements. We’ve demonstrated that inhibition of this process with the irreversible palmitoyl acyl-transferase (DHHC) inhibitor, 2-bromopalmitate (2BP), resulted in two changes that were distinguishable by dose and time properties (54). Short term treatments decreased SERT palmitoylation and reduced V_max_ to ∼60% without changes in surface or total SERT protein levels (54), indicating a role for palmitoylation in transport kinetics. At higher doses and longer treatment times, 2BP also caused a loss of SERT protein, suggesting that palmitoylation opposes transporter internalization and degradation (54). In the same study, we showed that treatment for 3 h with a therapeutic dose of escitalopram (500 nM) inhibited SERT palmitoylation and reduced SERT surface expression (54). These results were consistent with previous studies describing the internalization and loss of total SERT protein in the presence of chronic SSRI conditions both *in vitro* and *in viv*o (56–59).

Here we present our most recent investigation into uncovering the mechanism(s) behind the functional enhancement of SERT ASD coding variants, and how escitalopram may contribute to the rectification of these processes. Remarkably, we demonstrate enhanced palmitoylation for two variants (F465L and L550V), which was consistent with prior reports of enhanced V_max_ and surface expression for both variants (30). When challenged with either 2BP or escitalopram, F465L SERT palmitoylation was reduced, resulting in remediation of previously enhanced surface expression and transport capacity to basal WT levels. Collectively, these results highlight a novel component in the mechanism of functional enhancement for certain ASD associated hSERT coding variants, and may provide molecular evidence for the efficacy of SSRI therapeutics in the treatment of ASD.

## METHODS

### Cell Culture

Human Embryonic Kidney-293 (HEK293) cells stably expressing WT, C109A, I425L, F465L, L550V or K605N hSERT were grown in Dulbecco’s Modified Eagle Medium (DMEM) containing 5% fetal bovine serum, 100 μg/mL penicillin/streptomycin, and supplemented with 400 μg/mL of Geneticin (G418) for maintenance of stable expression. Cells were maintained in a humidified incubator gassed with 5% CO_2_ at 37°C. SERT expression level was verified by sodium dodecyl sulfate-polyacrylamide gel electrophoresis (SDS-PAGE) and immunoblotting of the cellular lysates against anti-human (h) SERT (Mabtechnologies-ST51-2) specific antibodies.

### Membrane Preparation

HEK293 cells expressing the indicated SERTs were grown in 100 mm poly-d-lysine coated plates to 85% confluency. Cells were washed twice with 3 mL of ice-cold Buffer B (0.25 mM sucrose, 10 mM triethanolamine, pH 7.8), scraped, and collected in 500 µL of Buffer B containing a protease inhibitor cocktail of 1 μM phenylmethylsulphonyl fluoride (PMSF) and 5 μM Ethylenediaminetetraacetic acid (EDTA) at 4°C and transferred to a 2 mL microcentrifuge tube. Cells were then pelleted *via* centrifugation at 3,000 *x g* for 5 min at 4^°^C, the supernatant fraction was removed, and the cell pellet was suspended in 1 mL of ice-cold Buffer C (0.25 M sucrose, 10 mM triethanolamine, 1 mM EDTA, 1 µM PMSF, pH 7.8) and subsequently homogenized *via* 30 strokes of the pestle in a Dounce homogenizer. Homogenates were cleared of cellular debris and nuclei by centrifugation at 800 *x g* for 10 min. The post-nuclear supernatant fraction was collected and centrifuged at 18,000 *x g* for 12 min at 4°C to pellet cell membranes. The resulting membrane pellet was suspended in 1mL of sucrose phosphate (SP) buffer (10 mM sodium phosphate, 0.32 M sucrose, pH 7.4 with 1 μM PMSF and 5 μM EDTA) and assayed for protein concentration.

### Acyl-Biotin Exchange (ABE)

The ABE method used as previously described (54) was adapted from Wan et. al. (60) where palmitoylated proteins are detected in three steps: (I) Free cysteine thiols are blocked; (II) thioester linked palmitoyl groups are removed by hydroxylamine (NH_2_OH); (III) the formerly palmitoylated and now newly generated sulfhydryl groups are biotinylated. Membranes prepared from HEK293 cells expressing the indicated hSERTs were solubilized in 250 μL of lysis buffer (50 mM HEPES pH 7.0, 2% SDS (w/v), 1 mM EDTA) containing 25 mM N-ethylmaleimide (NEM), incubated for 20 min in a 37°C water bath and mixed end-over-end for at least 1 hour at ambient temperature. Proteins were precipitated *via* the addition of 1 mL acetone and centrifugation at 18,000 *x g* for 10 min. The protein pellet was resuspended in 250 µl lysis buffer containing 25 mM NEM and incubated for 1 hour at ambient temperature. This process was repeated a final time with end-over-end mixing overnight at room temperature. NEM was removed *via* acetone precipitation/centrifugation and the protein pellet was resuspended in 250 μL 4SB buffer (50 mM Tris, 5 mM EDTA, 4% SDS, pH 7.4) and remnant NEM was removed by an additional acetone precipitation and centrifugation. The pellet was resuspended in 200 μL 4SB buffer and thioesterified palmitate molecules were removed by incubation with hydroxylamine (NH_2_OH); The sample was split into two equal aliquots (100 μL) with one diluted with 800 µl Tris-HCl, pH 8.0 (negative control) and the other diluted with 800 µL NH_2_OH (0.7 M final concentration) and incubated at ambient temperature for 30 min with end-over-end mixing. Both samples were treated with 100 μL of a sulfhydryl-specific biotinylating reagent, HPDP-biotin (0.4 mM final concentration) and incubated at ambient temperature with end-over-end mixing for 1 h. NH_2_OH and biotin reagents were removed by acetone protein precipitation with centrifugation at 18,000 *x g* and supernatant fraction aspiration. Protein pellets were then resuspended and solubilized in 150 μL 4SB, acetone precipitated with 600 µL, centrifuged at 18,000 *x g* followed with supernatant aspiration. The final pellet was suspended in 75 μL ABE lysis buffer. A 10 μL aliquot was set aside for determination of total SERT content by immunoblotting while 65 μL was diluted in 1500 μL Tris buffer and incubated with 50 μL of a 50% slurry of NeutrAvidin resin overnight at 4°C with end-over-end mixing. Unbound proteins were washed away by three cycles of 8,000 *x g* centrifugation, removal of the supernatant fraction, and resuspension in 750 μL radioimmunoprecipitation assay buffer (RIPA: 1% Tx-100, 1% sodium deoxycholate, 0.1% SDS, 125 mM sodium phosphate, 150 mM NaCl, 2 mM EDTA, 50 mM NaF). Proteins were eluted from the final pellet by incubation in 2x Laemmli sample buffer (SB: 125 mM Tris-HCl, 20% glycerol, 4% SDS, 200 mM DTT, 0.005% bromophenol blue) for 20 min at ambient temperature. Samples were then subjected to SDS-PAGE and immunoblotted for hSERT with ST51-2 antibody.

### Quantification of SERT palmitoylation by ABE

Samples (10 µl) taken from final resuspension (75 µl) just prior to NeutrAvidin extraction were used to directly assess total hSERT levels in the sample and normalize palmitoylation levels in this same sample. Band intensities were quantified using Quantity One software (Bio-Rad), normalized to total SERT protein present, and expressed as % control. Bands in immunoblots represent 13.3% of the total and 86.7% of the palmitoylated hSERT in each sample.

### Cell Surface Biotinylation

HEK293 cells stably expressing the indicated hSERT were grown in 24-well poly-d-lysine coated plates until 80% confluent. After treatments, the cells were washed three times with ice-cold Hank’s balanced salt solution containing Mg^2+^ and Ca^2+^ (HBSS Mg-Ca: 1 mM MgSO_4_, 0.1 mM CaCl_2_, pH 7.4), and subsequently incubated twice with 0.5 mg/mL of membrane-impermeable sulfo-NHS-SS-biotin for 25 min on ice with rocking. The biotinylation reagent was removed by aspiration and the reaction was quenched by two sequential incubations with 100 mM glycine in HBSS Mg-Ca for 20 min on ice with rocking. Cells were washed with HBSS Mg-Ca and then lysed with 250 μL per well RIPA buffer containing protease inhibitors. Lysates from 4 identically treated wells were pooled and assayed for protein content. Equal protein from each lysate pool (100 μg) was incubated with 50 µL of a 50% slurry of NeutrAvidin resin overnight at 4°C with end-over-end mixing. The protein bound resin was washed three times with RIPA buffer, and the bound protein was eluted with 32 μL of Laemmli SB followed by SDS-PAGE and immunoblotting for hSERT with ST-51-2 antibody. Equal amounts of total protein (10 μg) for each condition from the same samples were immunoblotted for total hSERT (ST51-2) in parallel with surface hSERT to account for differences observed in the genomic expression between WT and F465L hSERT and subsequent normalization of surface levels.

### Saturation analysis

HEK293 cells stably expressing WT or F465L hSERT were grown in 24-well poly-d-lysine coated plates until 80% confluent. The cells were washed twice with 0.5 mL 37°C Krebs Ringers HEPES (KRH: 120 mM NaCl, 5 mM KCl, 2 mM CaCl_2_, 1 mM MgCl_2_, 25 mM NaHCO_3_, 5.5 mM HEPES, 1 mM D-Glucose) buffer and 5HT uptake was conducted for 8 min at 37°C with 0.32, 0.6, 1, 3, 10, and 20 μM total 5HT containing 20 nM [^3^H]5HT; nonspecific uptake was determined in the presence of 100 nM (−)escitalopram. Cells were rapidly washed twice with 500 µL ice-cold KRH buffer and lysed with 1% Tx-100 for at least 20 min with rocking at ambient temperature. Radioactivity contained in lysates was assessed by liquid scintillation counting. Kinetic values were determined using Prism software, and V_max_ values were normalized to total cellular protein (pmol/min/mg) and transporter surface levels determined by surface biotinylation assays performed in parallel for each experiment.

### [^3^H]5HT uptake assay

HEK293 cells stably expressing WT or F465L hSERT were grown in 24-well poly-d-lysine coated plates until 80% confluent. The cells were washed with 0.5 mL 37°C KRH buffer and re-incubated in KRH buffer containing DMSO (control), 7.5 μM 2BP, or 500 nM escitalopram for 3 h at 37^0^C. The final DMSO concentration was 1% or less, which by itself did not affect 5HT transport activity. Immediately following incubation with the indicated treatments, cells were washed twice with 0.5 mL 37°C KRH buffer to remove treatment compounds and 5HT uptake was conducted for 8 min at 37°C in triplicate with 3 *μ*M 5HT (20 nM [^3^H]5HT) and nonspecific uptake was determined in the presence of 100 nM (−)escitalopram. Following uptake, the cells were rapidly washed 3 times with ice-cold KRH. Cells were then solubilized in 1% Triton X-100, and radioactivity contained in lysates was assessed by liquid scintillation counting.

### SDS-PAGE and Western Blotting

Proteins were denatured in Laemmli SB and were electrophoretically resolved using 4-20% polyacrylamide gels alongside a molecular weight protein standard. Proteins were then transferred onto polyvinylidene fluoride (PVDF) membrane and immunoblotted for hSERT using ST51-2 antibodies diluted 1:1,000 in blocking buffer (3% bovine serum albumin, phosphate buffered saline (PBS: 0.137 M NaCl, 0.0027 M KCl, 0.01 M NaH_2_PO_4_, 0.0018 M KH_2_PO_4_) and incubated for at least two h at ambient temperature or overnight at 4°C. After five washes for 5 min each with wash buffer (0.1% Tween 20, 1x PBS), the membrane was incubated for 45 min in alkaline phosphatase (AP)-linked anti-mouse IgG secondary antibody diluted 1:5,000 in blocking buffer followed by 5 additional 5 min washes. Protein bands were visualized by chemiluminescence using Immun-Star™ AP substrate (Bio-Rad) applied to the membrane and incubation for 5 min at ambient temperature. Band intensities were quantified using Quantity One® software (Bio-Rad).

## RESULTS

### Enhanced Palmitoylation of hSERT Coding Variants

Demonstrated in Figure 1A, a topographical model outlines hSERT’s 12 transmembrane domains (TMD) with potential intracellular cysteines for palmitoylation shown in large orange circles and five individual residues mutated in ASD associated hSERT coding variants shown in large green circles. The cysteines include: C15, C21, C147, C155, C357, C522, C540, C588, and C622. The coding variants associated with autism are designated as: G56A, C109A, I425L, F465L, L550V, and K605N. The three coding variants that confer an elevation in hSERT surface expression and 5HT transport are I425L, F465L, and L550V located in TMD’s 8, 9, and 11, respectively. Our previous work (54) indicates that hSERT palmitoylation supports increased 5-HT transport capacity and cell surface expression, leading us to investigate the potential role of palmitoylation in the enhancement of surface expression and transport capacity reported for autism-associated hSERT coding variants. Here, we subjected HEK293 cells stably expressing each coding variant to ABE for palmitoylation analysis.

**Fig. 1.**
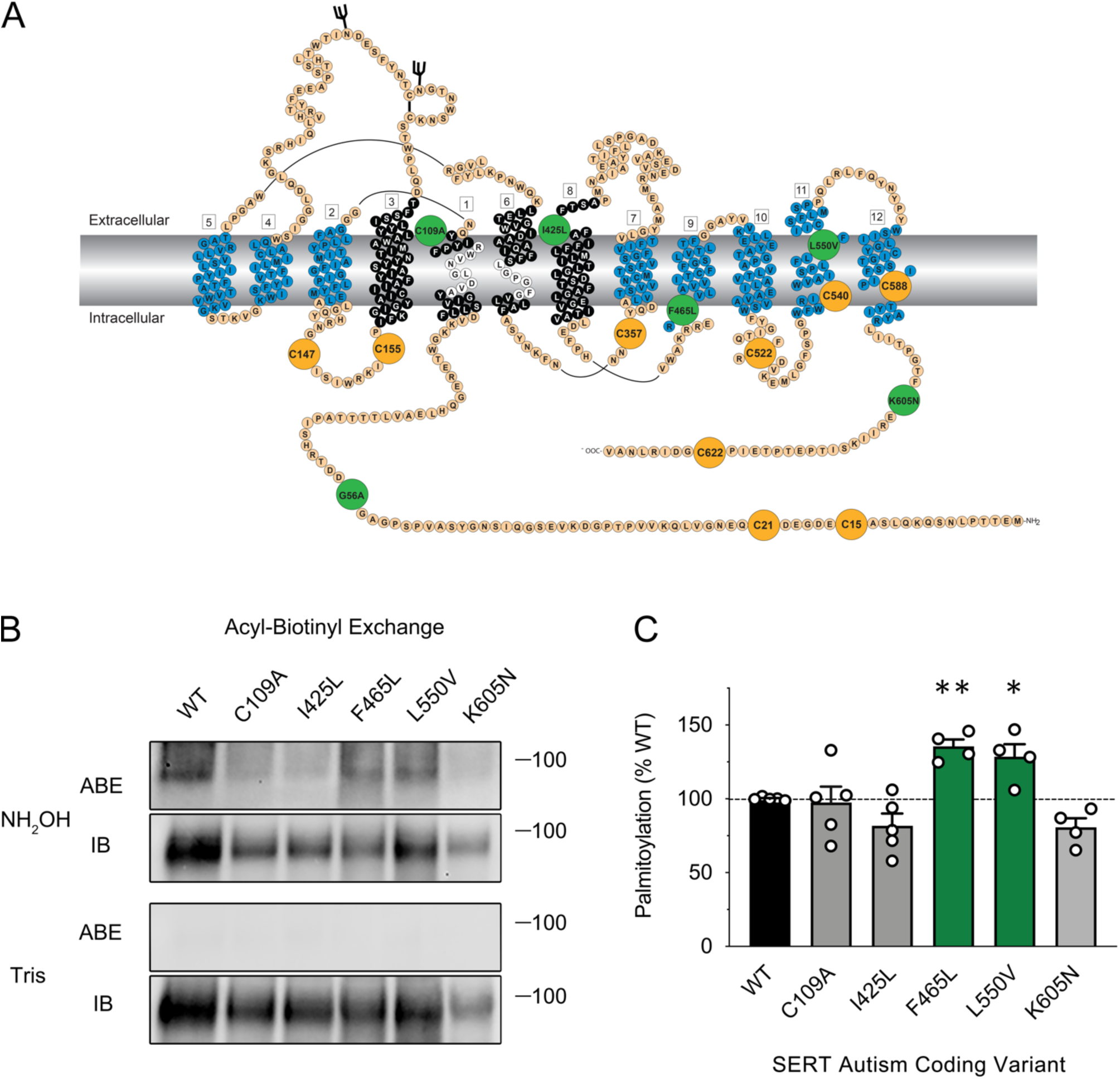
S-palmitoylation of hSERT autism spectrum disorder coding variants. (**A**) Topographical structure of the human serotonin transporter (hSERT) with 12 transmembrane domains. Intracellular cysteines that may serve as targets for palmitoylation are shown in enlarged orange circles and are designated: C15, C21, C147, C155, C357, C522, C540, C588, and C622. Coding variants identified in patients with autism are shown in large green circles and are designated: G56A, C109A, I425L, F465L, L550V, and K605N. (**B**) Indicated WT or mutant hSERT HEK293 cells underwent acyl-biotinyl exchange (ABE) and palmitoylation levels were analyzed by SDS-PAGE and immunoblotting (IB) with anti-hSERT antibody (ST51-2). (**C**) Quantification of palmitoylation normalized to total hSERT protein mean ± SEM of at least 4 independent experiments (n=4-5) relative to WT normalized to 100%. (** p < 0.01, * p < 0.05, one-way ANOVA with Fisher LSD post-test).

In these experiments, we utilized the ABE method as previously described (61–63) to analyze hSERT palmitoylation. In this method, palmitoyl-proteins are cleaved of their palmitoyl-moieties using hydroxylamine (NH_2_OH), followed with thioester modification by a sulfhydryl specific biotinylation reagent. The freshly biotinylated samples are subsequently pulled down *via* NeutrAvidin affinity chromatography to capture the previously palmitoylated and now newly biotinylated proteins. NeutrAvidin beads are then washed repeatedly, and the biotinylated proteins are eluted and subjected to SDS-PAGE and immunoblotted for hSERT (*Fig. 1B – NH_2_OH ABE*). Specificity of the assay for S-palmitoylation is determined *via* parallel assessment with Tris treatment rather than NH_2_OH where thioester-linked palmitoyl groups are not cleaved and there is no biotin conjugation and therefore no NeutrAvidin pull down (*Fig. 1B – Tris ABE*). Prior to incubation with NeutrAvidin, an aliquot of each sample is removed to determine total SERT levels in the sample post-ABE for normalization and quantification of palmitoylation (*Fig. 1B – NH_2_OH and Tris IB*).

Demonstrated in the ABE data and histogram of Figure 1B and C, SERT palmitoylation was increased in two hSERT coding variants relative to palmitoylation of the WT control group when normalized to their respective total hSERT protein levels post-ABE. F465L palmitoylation was increased to 135.5 ± 4.8% and L550V was increased to 128.5 ± 8.5% compared with WT (*Fig. 1C*, p<0.01 and p<0.05, respectively, *via* ANOVA with Fisher LSD post-test samples, n=4). In some cases, palmitoylation was unchanged including coding variants C109A (*Fig. 1C,* 97.4 ± 10.9% *via* ANOVA with Fisher LSD post-test samples, n=5) and I425L (*Fig. 1C*, 81.7 ± 8.4% *via* ANOVA with Fisher LSD post-test samples, n=5). This revealed to us that two of the three TM located coding variants (F465L and L550V) previously associated with increased surface expression and transport capacity also maintained an elevated state-of-palmitoylation compared to the WT transporter.

### Enhancement of F465L hSERT Surface Expression and Transport Capacity

Before we pursued a deeper analysis of the mechanism underlying the enhanced coding variants, we elected to validate the previously demonstrated enhancement of surface and transport capacity in F465L hSERT (30). For the purposes of our study, we examined F465L hSERT alone, which demonstrated greater statistical significance in our initial palmitoylation analysis compared to L550V hSERT. In contrast to previous studies on F465L hSERT, our studies were conducted in HEK293 cells stably expressing hSERTs and not in transiently transfected HeLa cells (30). Our initial ABE analysis of F465L hSERT suggested there may be a reduction in total F465L expression compared to WT (*Fig. 1*). To further analyze this, we ran equivalent protein amounts from cellular lysate of HEK293 cells expressing WT or F465L hSERT on SDS-PAGE and immunoblotted for hSERT. Demonstrated in Figure 2A, the total expression of F465L was 47.5 ± 6.9% compared to WT in our stably expressing HEK293 cell lines (p<0.001 *via* Student’s t-test, n=4). This was not surprising since stable transformants can express different levels of the target protein based on the site of cDNA incorporation into the cellular genome. To more accurately compare the differences between WT and F465L hSERT surface expression, we assessed surface expression by normalizing our cell surface values to an even level of total hSERT by dividing the density of surface hSERT by its respective total hSERT level. When this was performed, we observed a statistical increase in the surface density of F465L by 124.9 ± 9.1% compared with WT (*Fig. 2B*, p<0.05 *via* Student’s t-test, n=9). We then determined 5HT transport capacity for the WT and F465L expressing cells and normalized values to cellular protein levels (pmol/min/mg) and cell surface density determined in parallel with equal protein loads. With WT normalized to 100%, we observed an increase in F465L transport capacity to 129.5 ± 4.1% of WT control (*Fig. 2C*, p<0.01 *via* Student’s t-test, n=3). Together, these data indicate that F465L displays enhanced transport capacity and cell surface expression as previously described, and correlates enhanced transport and surface expression with enhancement of F465L palmitoylation (*Fig. 1B and C*).

**Fig. 2.**
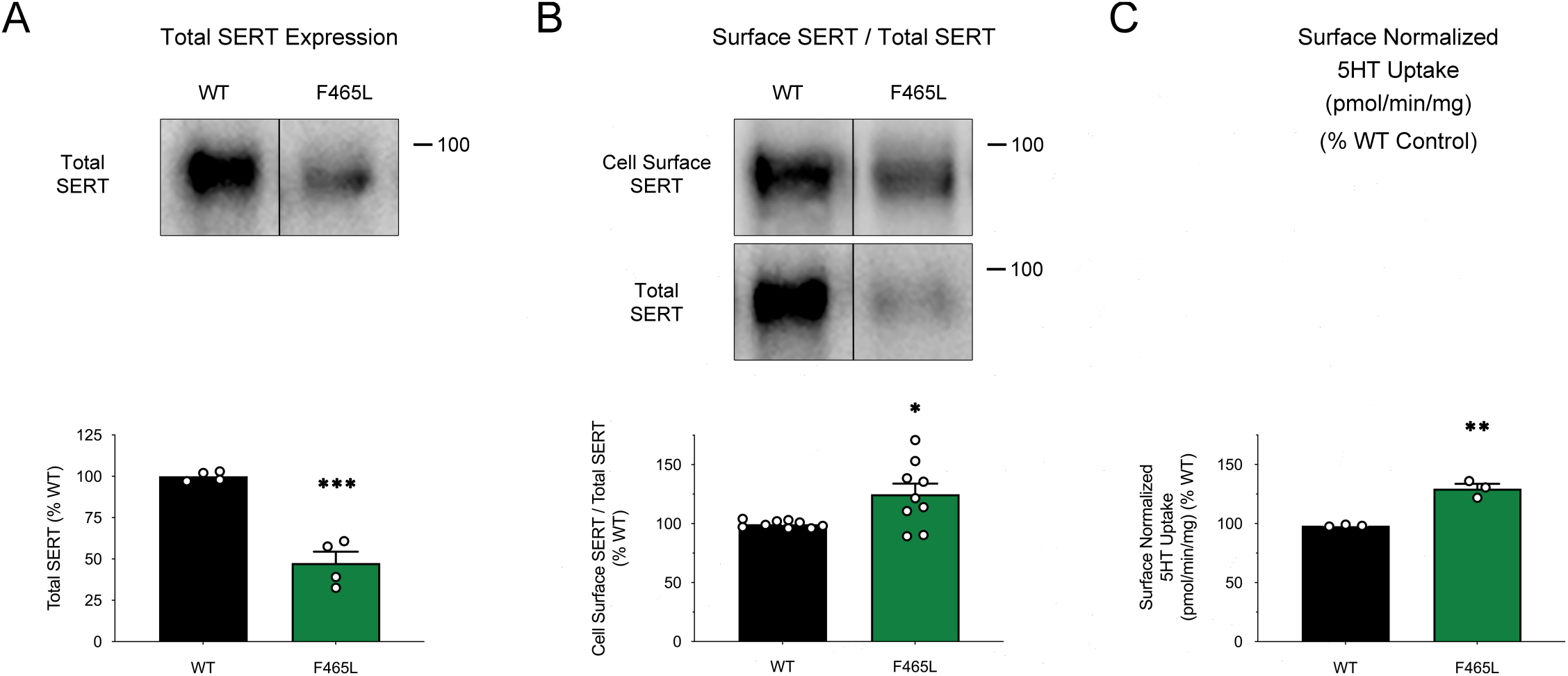
Enhancement of F465L surface expression and transport capacity. HEK293 cells expressing WT or F465L hSERT were analyzed for (**A**) total hSERT protein levels when equivalent cellular protein was loaded onto SDS-PAGE followed with immunoblotting by anti-hSERT (ST51-2). Line within the box (|) indicates splicing from the same gel. Data presented represent mean ± SEM of 4 independent experiments (*** p<0.001 relative to WT control, Student’s t-test for independent samples). (**B**) WT or F465L hSERT HEK293 cells underwent cell surface biotinylation and hSERT surface expression was analyzed by SDS-PAGE and immunoblotting. Total hSERT was also assessed in parallel with equal cellular protein loads (10 µg) to account for the decrease in stable expression between WT and F465L hSERT. Line within the box (|) indicates splicing from the same gel. Mean ± SEM of 9 independent experiments (n=9) relative to WT normalized to 100%. (* p < 0.05 versus WT control, Student’s t-test for independent samples). (**C**) 5HT transport was performed and normalized to total cellular protein (pmol/min/mg) and cell surface levels. Mean pmol/min/mg ± SEM of 3 independent experiments performed in triplicate relative to WT normalized to 100%. (** p<0.01 relative to WT control, Student’s t-test for independent samples).

### F465L hSERT Demonstrates Enhanced Transport Kinetics

Our next step was to investigate if enhanced palmitoylation was consistent with enhanced 5HT transport kinetics (k_cat_) or entirely the result of increased cell surface expression. Previous studies on F465L kinetics revealed an increase in V_max_ with lowered K_m_ values compared to WT control (30). We have demonstrated hSERT V_max_ is dependent on hSERTs state-of-palmitoylation, with acute inhibition (30 min) by 2BP decreasing hSERT V_max_ without changing K_m_ or surface expression patterns. To investigate this, we conducted 5HT uptake saturation analysis with WT and F465L hSERT with transport activity normalized to total cell protein levels (pmol/min/mg) and cell surface expression of hSERT determined in parallel. The results from these studies demonstrated that surface normalized V_max_ was consistent with palmitoylation status, increasing to 156.0 ± 20.8% of the WT control (*Fig. 3A, B*, p<0.05 *via* Student’s t-test, n=4). These data also revealed no change in the K_m_ of F465L compared to WT (*Fig. 3C*, *via* Student’s t-test, n=3). Demonstrated in Table 1, V_max_ values for WT and F465L were 61.1 ± 4.4 pmol/min/mg and 97.2 ± 9.1 pmol/min/mg, respectively (p<0.05 *via* Student’s t-test, n=4) without any change in K_m_. These data were consistent with previously published data on F465L demonstrating a kinetic enhancement that was independent of surface expression (30) where F465L V_max_ was elevated by 56.0% consistent with Prasad and colleagues previous work (30) (*Fig. 3A and B*, green saturation curve, p<0.05 *via* Student’s t-test, n=4)

**Fig. 3.**
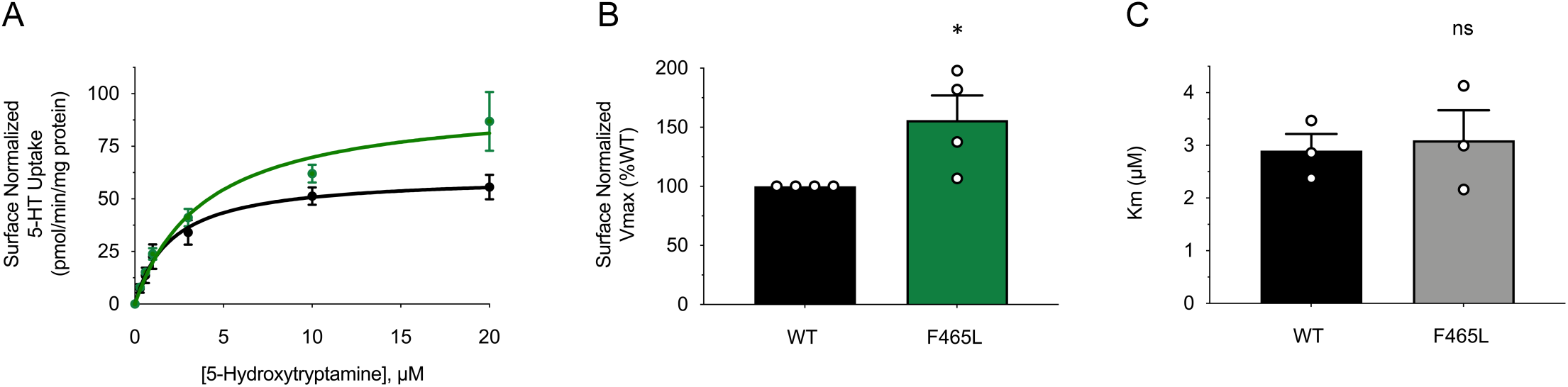
F465L hSERT demonstrates enhanced V_max_ without changes in K_m_. HEK293 cells expressing WT or F465L hSERT were subjected to 5HT transport saturation analysis (**A**) Saturation analysis was performed and normalized to total cellular protein (pmol/min/mg) and cell surface levels. Data presented represents mean ± SEM of 4 independent experiments normalized to surface hSERT and fit to Michaelis-Menten kinetics. (**B**) Data are presented as means of percentage V_max_ ± SEM of three independent experiments performed in triplicate. (* p<0.05 relative to WT control, Student’s t-test for independent samples). (**C**) Quantification of K_m_ values are presented as means of K_m_ ± SEM of at least 3 independent experiments relative to control (ns indicates no significance, Student’s t-test for independent samples).

**Table 1.**
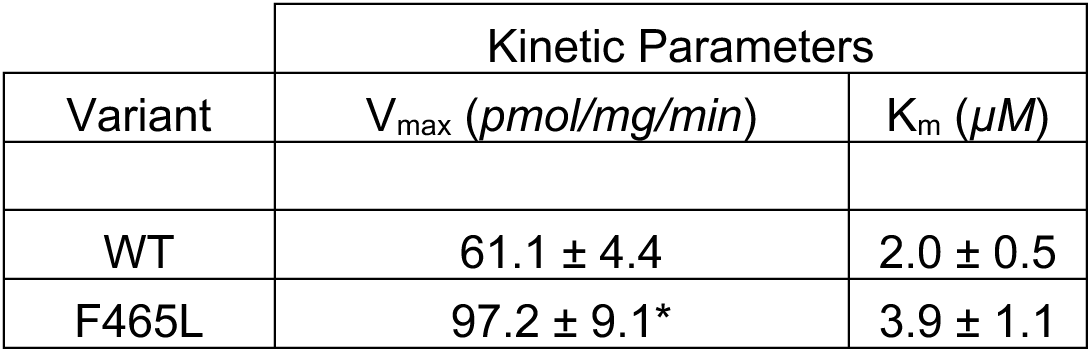
Kinetic parameters of autism-associated F465L hSERT. WT or F465L hSERT HEK293 cells were subjected to 5HT transport saturation analysis. Data are presented as means of percentage V_max_ or K_m_ ± SEM of three independent experiments performed in triplicate. (* p<0.05 relative to WT, Students t-test for independent samples compared to WT control).

| Variant | Kinetic Parameters |  |
| --- | --- | --- |
| | $V_{\max}$ (pmol/mg/min) | $K_m$ ($\mu M$) |
| WT | 61.1 ± 4.4 | 2.0 ± 0.5 |
| F465L | 97.2 ± 9.1* | 3.9 ± 1.1 |

### Escitalopram and 2BP Diminish Enhanced F465L hSERT Palmitoylation

Our previous work with hSERT has shown that palmitoylation is sensitive to inhibition with 500 nM escitalopram (30). Within 3 h of treatment, we demonstrated that escitalopram reduces hSERT palmitoylation consistent with a corresponding reduction in hSERT surface expression. Because of this, we wondered if/how escitalopram may impact our new findings demonstrating enhanced F465L hSERT palmitoylation, surface expression, and transport capacity. To answer this question we treated HEK293 cells expressing either WT or F465L hSERT with 7.5 µM 2BP as a positive control or 500 nM escitalopram followed by palmitoylation analysis by ABE, and is consistent with concentrations used in our prior study on modulation of SERT by escitalopram (54).

To contend with the notable differences in stable expression, we determined hSERT protein levels post-ABE and loaded equal SERT onto the neutravidin resin and total hSERT immunoblot for ABE normalization (*Fig. 4A - IB).* In at least four independent experiments (n=4-11), we found that 2BP decreased WT hSERT palmitoylation to 65 ± 7% of WT vehicle control (*Fig. 4B*, p<0.05 *via* ANOVA with Tukey post-test, n=4), consistent with our previous findings in LLC-PK_1_ cells expressing HA-hSERT (30). When treated with 500 nM escitalopram, palmitoylation of hSERT was significantly decreased to 56.0 ± 9.7% of WT vehicle control (*Fig. 4B*, p<0.01 *via* ANOVA with Tukey post-test, n=4). This result mirrored our findings with 2BP treatment and was consistent with our previous data (30).

**Fig. 4.**
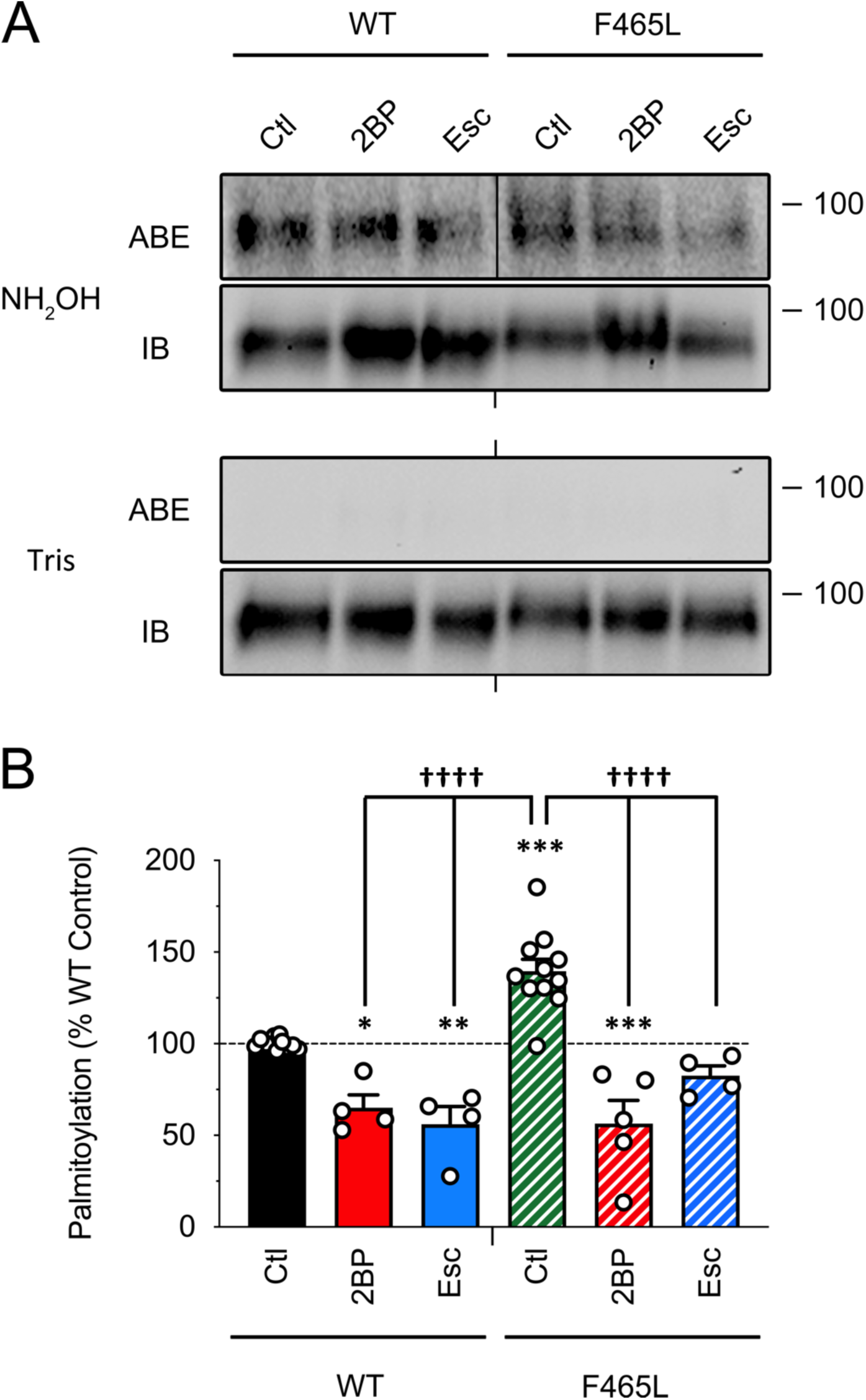
Escitalopram and 2BP decrease previously enhanced F465L palmitoylation. HEK293 cells expressing WT or F465L hSERT were treated with 7.5 µM 2BP or 500 nM escitalopram (Esc) for 3 h followed with assessment of palmitoylation levels using ABE. Line within the box (|) indicates splicing from the same gel. (**A**) Western blot representative of at least four independent experiments (n = 4-11) for WT or F465L hSERT under control (Ctl), 2BP, or Esc treatment conditions. (**B**) Quantification of palmitoylation, mean ± SEM of at least four independent experiments (n = 4-11) performed relative to WT control normalized to 100%. (* p<0.05, ** p<0.01, ***, p<0.001 versus control, ^††††^ p < 0.0001 between indicated treatment conditions, one-way ANOVA with Tukey post-test).

We next determined the impact of 2BP and escitalopram on F465L palmitoylation. Here, palmitoylation analysis of F465L hSERT by ABE again revealed a statistically significant increase in palmitoylation of 139.6 ± 6.5% compared to WT control (*Fig. 4B*, p<0.001 *via* ANOVA with Tukey post-test, n=11). We then treated F465L hSERT with 7.5 µM 2BP and observed a 43.7 ± 12.7% decrease from WT vehicle control and 83.3 ± 12.7% reduction from F465L vehicle control (*Fig. 4B*, p<0.001 and p<0.0001, respectively *via* ANOVA with Tukey post-test, n=5). This finding suggested to us that F465L may configure hSERT into a conformation that increases its sensitivity to palmitoylation. When F465L was treated with 500 nM escitalopram, we saw a 57.0 ± 5.4% decrease in palmitoylation from F465L vehicle control (*Fig. 4B*, p<0.0001 via ANOVA with Tukey post-test, n=4) that was not statistically different from WT vehicle control (*Fig. 4B*, 82.6 ± 5.4% *via* ANOVA with Tukey post-test, n=4). Remarkably, this revealed that escitalopram-treated F465L hSERT palmitoylation was reduced to basal WT levels when compared to the WT hSERT control. This suggests to us that escitalopram therapeutic strategies in ASD patients may function by decreasing a hyper-palmitoylated mutant transporter to basal WT palmitoylation levels in addition to directly inhibiting 5HT transport.

### Remedial impact of Escitalopram and 2BP on F465L hSERT Surface Expression and Transport Capacity

The results of our palmitoylation data led us to believe that the hyper-functional characteristics of F465L could be remitted with escitalopram. Because we have shown that stimulating hSERT palmitoylation by overexpression of DHHC enzymes (*unpublished data*) leads to enhanced hSERT surface expression and 5HT transport capacity, coupled with our previous findings that these processes are sensitive to inhibition with 2BP, we proceeded to test if this coding-variant induced enhancements were sensitive to downregulation.

To test if these processes were consistent with the changes observed with palmitoylation, we tested hSERT surface expression after being challenged with 7.5 µM 2BP or 500 nM escitalopram for 3 h. Demonstrated in Figure 5A, we observed an inhibition of WT hSERT surface expression when treated with either 2BP (80.1 ± 4%) or escitalopram (72.2 ± 4.4%) (*Fig. 5B*, p<0.05 and p<0.01, respectively compared to control via ANOVA with Fisher LSD post-test, n=5-6). When compared with WT control surface expression, untreated F465L hSERT was enhanced to 124.9 ± 9.1% (*Fig. 5B*, p<0.05 via ANOVA with Fisher LSD post-test, n=9). When treated with 2BP or escitalopram, we saw a reduction in F465L hSERT surface expression that was returned to baseline compared with WT hSERT, suggesting that inhibition of palmitoylation by these processes leads to a reduction in their surface expression (*Fig. 5B*, p<0.05 vs F465L via ANOVA with Fisher LSD post-test, n=4-6).

**Fig. 5.**
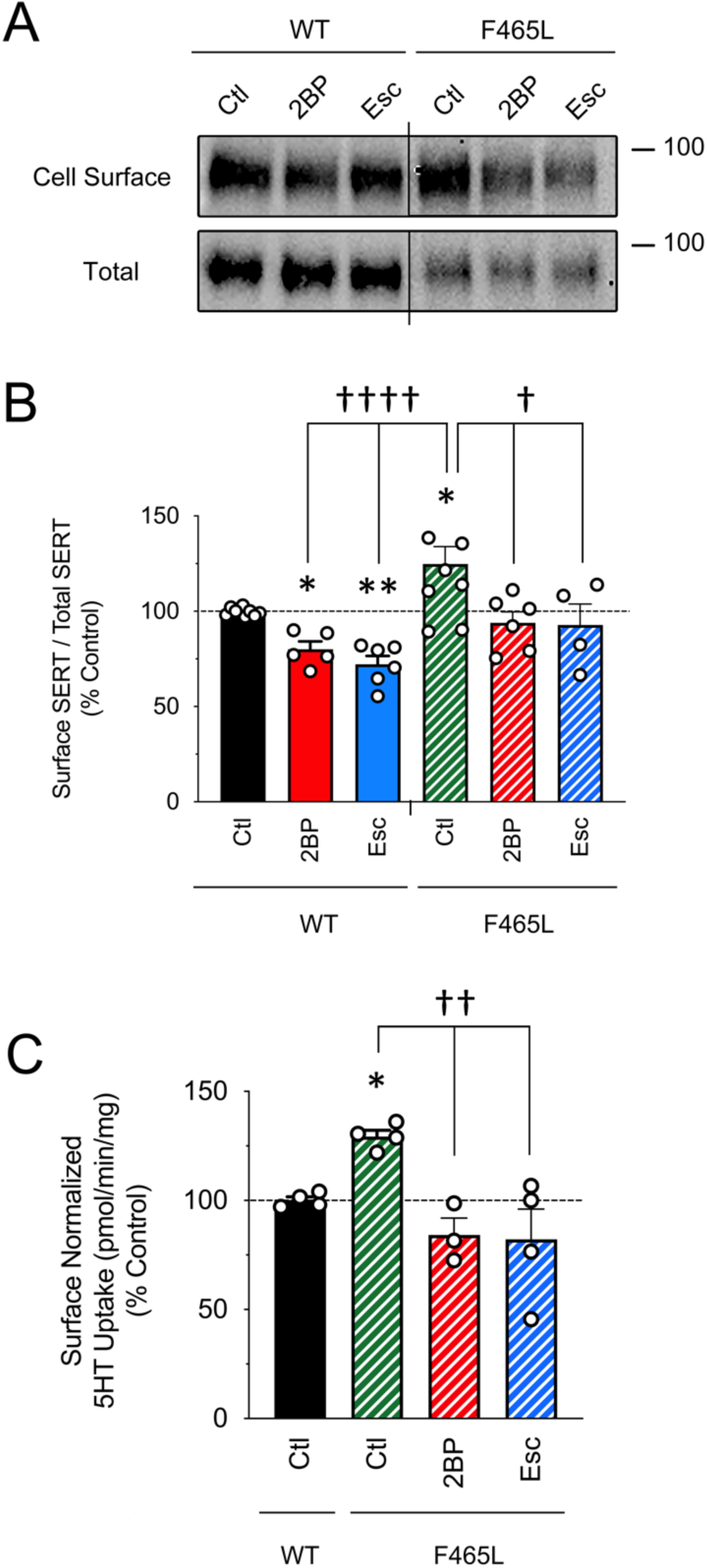
Escitalopram and 2BP decrease previously enhanced F465L surface expression and transport capacity. HEK293 cells expressing WT or F465L hSERT were treated with 7.5 µM 2BP or 500 nM escitalopram (Esc) for 3 h followed by assessment for SERT surface expression. (**A**) Western blot representative of WT or F465L hSERT surface levels under control (Ctl), 2BP, or Esc treatment conditions in at least four independent experiments (n = 4-7). (**B**) Quantification of hSERT cell surface blots normalized to total SERT levels, mean ± SEM of at least four independent experiments (n = 4-7) relative to WT control normalized to 100%. (* p<0.05, *p<0.01 condition vs WT Ctl, ^†^ p<0.05, ^††††^ p<0.0001 vs indicated conditions, one-way ANOVA with Fisher LSD post-test). (**C**) Quantification of 5HT uptake, mean ± SEM of four independent experiments (n = 4) performed relative to WT Ctl normalized to 100%. (* p<0.05 vs WT Ctl, ^††^ p<0.01 vs F465L Ctl, one-way ANOVA with Fisher LSD post-test).

Likewise, we next assessed the transport capacity of F465L against WT hSERT under control, 2BP, and escitalopram treatment conditions. When corrected for surface levels (as previously described), we found that F465L transport was enhanced (136 ± 4.1%) compared to WT control (*Fig. 5C*, p<0.05 via ANOVA with Fisher LSD post-test, n=4). When F465L was treated with 2BP, 5HT transport was reduced to baseline, at 84.2 ± 7.7% compared to WT control (*Fig. 5C*, p<0.01 compared to F465L control via ANOVA with Fisher LSD post-test, n=4), reflecting the changes we observed in our palmitoylation and surface analyses. Likewise, escitalopram reduced F465L transport to 82.2 ± 13.8% compared to WT control (*Fig. 5C*, p<0.01 compared to F465L control via ANOVA with Fisher LSD post-test, n=4), again consistent with changes we observed in our earlier palmitoylation and surface studies. It is prudent to mention again, that changes in transport were assessed following diligent and vigorous washing to remove and dilute the original treatment-enriched media, before the addition of substrate. Ultimately, these data confirm previous studies that F465L surface expression and transport is elevated compared to WT hSERT and implicate increased palmitoylation in the F465L hSERT variant as a mechanistic component for enhanced function, with an apparent remedial impact via inhibition with 2BP or escitalopram.

## DISCUSSION

In the current study, we investigated the mechanism behind the enhanced functional phenotype (elevated surface expression and transport capacity) of rare autism-associated hSERT coding variants. Additionally, we reveal a potential mechanism behind the reported efficacy of SSRIs as a therapeutic approach for ASD patients. Our previous work, and that of Harada et. al, confirmed that hSERT is a palmitoyl-protein (53–55), adding a layer to the increasingly complex mechanism underlying serotonergic homeostasis. We’ve demonstrated that palmitoylation of hSERT drives three major functional consequences, including enhanced kinetic activity, elevated surface expression, and increased total hSERT protein levels (54). When palmitoylation is inhibited for 30 min, SERT V_max_ is reduced without changes in K_m_, surface expression, or total hSERT levels. When treated with higher 2BP concentrations in longer time intervals, we observed losses in hSERT surface expression and total hSERT density that were consistent with losses in palmitoylation (54). In unpublished data, we have observed an enhancement in hSERT surface expression following promotion of hSERT palmitoylation by DHHC overexpression, further contributing to the dynamic hypothesis of hSERT membrane localization dependent on palmitoylation status. We have also previously revealed that escitalopram-induced internalization of hSERT (57, 64) parallels palmitoylation status (54–56, 65), and that this process was not reversible in wash-out conditions contrasting prior reports (54, 65). These previous findings implicate palmitoylation as a control mechanism for hSERT kinetics, surface retainment, and maintenance of total protein levels. Furthermore, treatment with escitalopram modulates hSERT palmitoylation, promoting dramatic losses in surface hSERT, total hSERT protein, and SERT mRNA expression levels in immortalized serotonergic neurons and the Raphe nuclei (58, 59, 66).

The rare gain-of-function hSERT mutations found in patients with autism are classified by their location in the hSERT amino-acid chain and their phenotypic enhancement. The G56A and K605N mutants are located on the intracellular N- and C-termini respectively, and have been reported to enhance hSERT transport capacity without changing SERT trafficking or total expression (33, 35). These mutants also display an insensitivity to PKC, PKG, and p38 MAPK signaling pathways, which normally maintain hSERT surface expression and transport capacity through surface recruitment (PKG and p38 MAPK) and/or endocytosis (PKC) (29, 37, 46–48, 67–69). The ASD-associated hSERT mutations I425L, F465L, and L550V are located within the transmembrane domain regions and exhibit enhanced surface expression and transport capacity (30). The F465L and L550V hSERT mutations exist in the 9^th^ and 11^th^ transmembrane domains, respectively (Figure 1), which are recognized to be highly conserved across all SLC6 family members (30). Prior investigators hypothesized that these functional outcomes may be the result of newly stabilized conformations that promote recruitment of hSERT to the cellular surface (30). It has been demonstrated that two of the ASD-associated hSERT mutants, G56A and K605N, induce a conformational change that support an open-outward 5HT binding conformation *in vitro* and *in vivo* (70). In addition, SERT conformational status has been found to dictate its state of phosphorylation (71), implying that coding variant induced conformation changes may modulate its PTM status. This information led us to consider that the enhanced SERT surface expression demonstrated by I425L, F465L, and L550V may be the result of a mutation-induced conformational change that promotes enhanced palmitoylation, perhaps by promoting DHHC-hSERT interactions or protein-hSERT interactions that are controlled by hSERTs palmitoylation status. Additionally, because prior clinical reports have outlined the efficacy for SSRIs on modulating autism-induced anxiety and abnormal brain activation (40–45), we hypothesized that escitalopram would decrease hSERT palmitoylation and normalize function, as we have seen previously (54).

For the purposes of our study, we were kindly gifted previously established HEK293 cells stably expressing each coding variant and subjected them to ABE analysis (Figure 1). The primary challenge in using these stable cell lines was the inconsistent levels of expression between each coding variant, which required us to use normalization strategies for subsequent surface and transport analysis. The ABE assay already utilizes an internal control where palmitoylation is normalized to the respective total protein of interest, with palmitoylation quantified as a fraction of the respective level of total transporter exposed to the NeutrAvidin resin. Our analysis revealed two hSERT variants (F465L and L550V) with elevated palmitoylation compared to WT hSERT (Figure 1), which was consistent with palmitoylation mediated elevation of surface expression and transport capacity (54, 55). Following our palmitoylation analysis, we focused our functional characterization on the F465L variant where we confirmed functional enhancement of surface expression and transport capacity as previously reported (30). In addition, we conducted 5HT transport saturation analysis and normalized the transport values to surface SERT expression to determine if the enhanced transport capacity of the F465L variant was due exclusively to increased surface expression or enhanced transport kinetics. Results from these experiments revealed an increase in V_max_ with no change in K_m_. Interestingly, when F465L hSERT transport capacity was normalized to total cellular protein (pmol/min/mg) and surface expression, V_max_ remained enhanced suggesting that the catalytic nature of F465L is independent of surface density. This data is consistent with our previous findings, demonstrating that palmitoylation controls hSERT (54) and dopamine transporter (DAT) (61–63) V_max_ without changing K_m_, suggesting that this mutation may positively drive palmitoylation and secondarily enhance its kinetic properties.

The evidence outlining the enhancement of hSERT surface recruitment was highly suggestive to us that a post-translational event may be driving hSERT to the surface as we and Harada et. al have seen previously (54, 55). Although coding-variant induced changes in enzyme kinetics and substrate recognition is well-accepted (71), it seems unlikely that an amino-acid substitution would be solely responsible for changes in trafficking patterns without promoting changes in co- or post-translational processing. It is possible that mutational alteration of hSERTs tertiary structure by the ASD coding variants may promote receptivity of hSERT intracellular cysteine residues for palmitoylation by DHHC enzymes. Harada et. al suggest that C147 and C155 (Figure 1), found within the first intracellular loop, are palmitoylated (55). In addition, it has been reported via mass-spectrometry that C622 (Figure 1), found within the C-terminus, is palmitoylated as well (72). We have observed that enhancing palmitoylation through differential DHHC over-expression promotes hSERT recruitment to the cellular surface with independent changes in hSERT kinetic properties (unpublished data), exemplifying a potential role for differential impacts by DHHC enzymes. This highlights the possibility that independent DHHC enzymes, found in different gross tissues and subcellular compartments, may be inducible by unique physiologic signals to enhance palmitoylation of hSERT with different functional outcomes. For example, activating DHHCs located within the plasma membrane may function hyper-acutely by raising hSERT V_max_ independent of surface levels, while vesicular-located DHHCs may function by acutely raising hSERT surface expression, and Golgi and/or ER DHHCs may function sub-acutely/chronically by promoting the biogenesis and maturation of hSERT. We have observed this phenomenon in our investigations of the human norepinephrine transporter (hNET), with ER-located DHHC1 apparently driving total and surface hNET expression, without coexisting changes in intrinsic hNET kinetic properties (73). Likewise, with DAT we have observed palmitoylation modulates DAT kinetics and total expression, without alterations in surface patterns (61, 62). All three monoamine transporters (MAT) are unique in this fashion, with hNET having four (73), DAT having five (62), and hSERT having nine available intracellular cysteines (Figure 1), implying that hSERT is capable of more diverse functional outcomes via site-directed palmitoylation compared to hNET and DAT. Ultimately, it would be prudent to discern the functional implications of each palmitoylated cysteine residue, publish our observations on the functional impacts following differential DHHC overexpression, and characterize the potential structure-mediated changes among disease variants that facilitate changes in SERT PTMs and subsequent functional consequences.

In our therapeutic experiments, escitalopram reduced F465L palmitoylation to the basal levels reflected by WT control (Figure 4), suggesting that the mechanism of escitalopram’s efficacy as an ASD therapeutic may be due to its modest inhibition of hyper-palmitoylated hSERT to a basal WT state. Likewise, when SERT palmitoylation was inhibited by 2BP or escitalopram, hSERT surface expression and transport capacity were reduced to basal WT levels (Figure 5). Represented by our working model in Figure 6, we propose that the mechanism underlying escitalopram’s efficacy in this scenario may be due to reducing hSERT palmitoylation, which then decreases hSERT surface levels and subsequent transport capacity. Rigorous discussion on this mechanism has brought to our attention the important aspect of initial and sustained orthosteric inhibition of hSERT by escitalopram. In other words, it was believed that if escitalopram is already providing inhibition of hSERT uptake to a nearly maximum degree, then what is it’s therapeutic utility in modulating hSERT palmitoylation and functional outcomes? The possibility for this novel mechanism partially relies on data published regarding hSERT occupancy studies with escitalopram. A recent meta-analysis (74) has organized the data from six independent studies on hSERT occupancy with escitalopram across different dosages and brain regions following one day of intake. Collectively, these studies revealed that hSERT occupancy with escitalopram varies widely, with 5 mg equating to 50 ± 1 – 67 ± 7% occupancy (75, 76), 10 mg equating to 59 ± 5% occupancy (75–80), 20 mg equating to 65% - 81% occupancy (75, 76, 78), and 30 mg equating to 62 ± 3 – 79 ± 6% occupancy (76). These data reveal that hSERT occupancy trends in a dose-dependent manner, and that a standard clinical starting dose of escitalopram (5-10 mg) often leads to slightly greater than half occupancy of available hSERT’s. This implies a substantial portion of hSERT is not occupied by escitalopram at therapeutic levels, allowing for continued global serotonin transport. In the context of the hyper-palmitoylated, hyper-functional hSERT ASD mutants, it seems possible that treatment with a therapeutic dose of escitalopram would bind 50-60% of available SERTs, leading to reduced palmitoylation and thus internalization of a subset of these transporters. This mechanism effectively removes a portion of hSERT from the available pool of physiologically active transporter at the cellular surface, allowing the remaining unbound pool of functionally enhanced hSERT to operate at a reduced capacity via decreased surface density, establishing a new equilibrium and returning global serotonergic tone to basal levels (Figure 6). However, the mechanisms underlying serotonergic terminal remodeling in health and disease are unclear, and highly complex, likely involving both immediate effects from acute hSERT blockade and also chronic changes in serotonergic machinery, deserving further exploration and study.

**Fig. 6.**
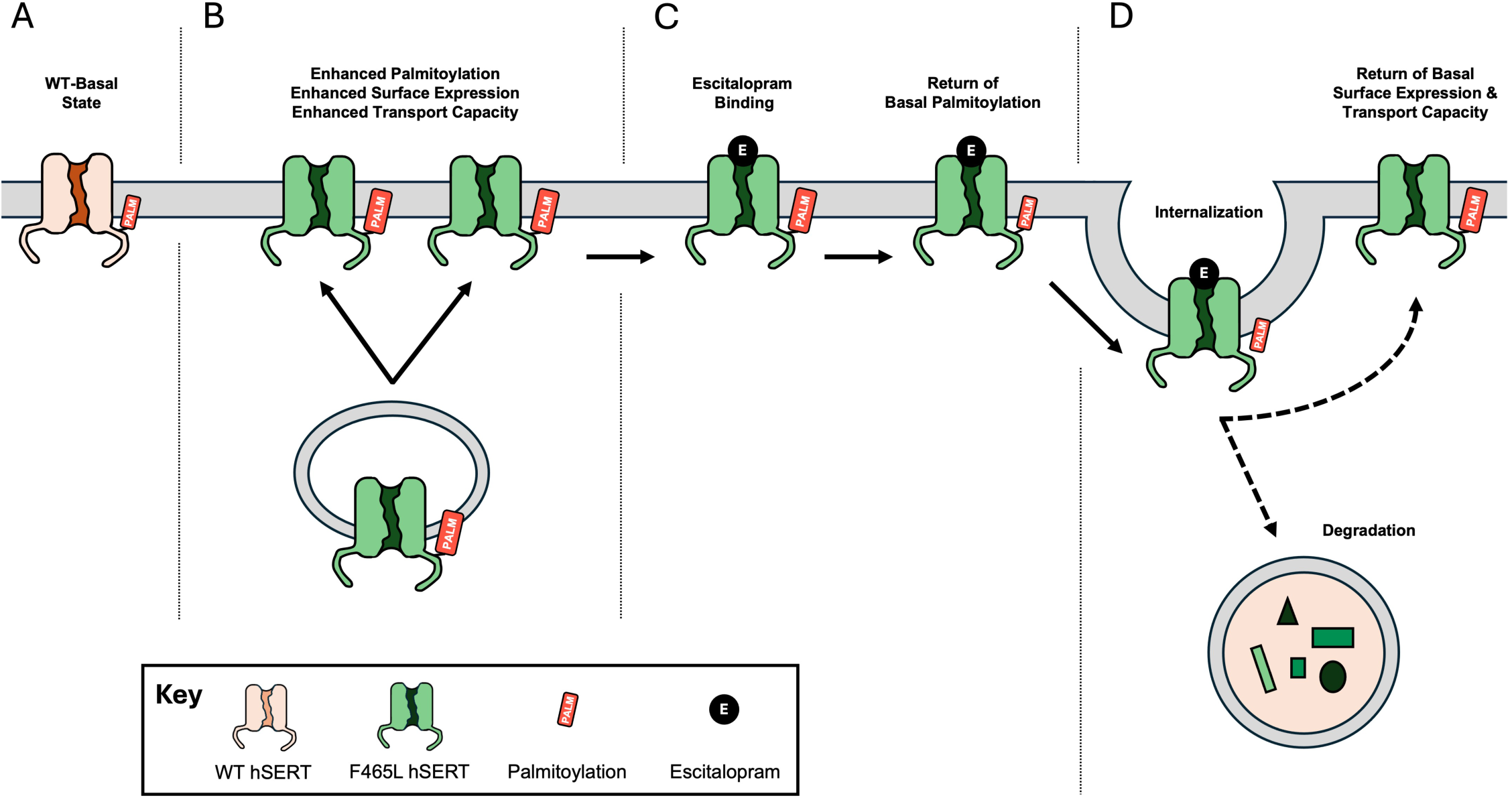
Proposed pathogenic-to-therapeutic model for the role of SERT palmitoylation in escitalopram modulated autism spectrum disorder. (**A**) WT hSERT under basal conditions is regulated dynamically by palmitoylation controlling kinetics, trafficking, and protein expression patterns in accordance with physiologic requirements. (**B**) F465L (and L550V) hSERT demonstrate increased palmitoylation compared to WT hSERT, potentially via induced-conformations that may increase accessibility of cysteine residues for palmitoylation. Enhanced palmitoylation of F465L manifests in increased surface expression and 5HT transport capacity compared to WT hSERT. (**C**) Therapeutic doses of escitalopram bind SERT (at an unknown site), potentially locking SERT in a different conformation that promotes a decrease in SERT palmitoylation - either through inhibition of palmitoyl-attachment, promotion of palmitoyl-release, or both. (**D**) Decreasing F465L palmitoylation (via 2BP or escitalopram) leads to decreased SERT surface expression, presumably through SERT endocytosis (described as *internalization*) and/or disrupted surface recruitment, with a subsequent reduction in 5HT transport capacity to basal WT levels (dashed lines).

Defined by the conformation of TMs 1b, 6a, 10, 11, and EL4 and EL6 (81), the allosteric site of hSERT is a secondary binding site which is known to modulate hSERT function (81–83). Recent hypotheses have speculated that escitalopram’s effectiveness may be dependent on binding of the allosteric site, and not simply via inhibition of the orthosteric site (58, 83–86). Two mechanisms on this have been postulated. Firstly, it has been demonstrated that escitalopram binding to the allosteric site directly obstructs orthosteric site dissociation, further enhancing extracellular 5HT levels (81–83). However, it has also been proposed that escitalopram binding to the allosteric site can influence the physical interaction between SERT and its interacting proteins (83). These observations imply that escitalopram (or 5HT) binding to the allosteric site can modulate the properties of the orthosteric binding site through more than one mechanism, effectively operating as an allosteric serotonin reuptake inhibitor (81–83). Interestingly, 5HT itself causes acute down-regulation of hSERT surface expression and surface normalized V_max_ (87), directly mirroring our findings following inhibition of hSERT palmitoylation (54), suggesting an allosteric effect from 5HT (or SSRI) binding that may be due to induced changes in palmitoylation. While the mechanisms underlying escitalopram-induced changes in hSERT trafficking are not fully understood, one study has reported them to be independent of allosteric site binding (58). However, this conclusion was established following site-directed mutagenesis of five amino acids in TM 10 and 12 - A505V, L506F, I507L, S574T, and I575T - effectively disabling escitalopram binding to the allosteric site (58, 83, 88, 89). Furthermore, the authors concluded that these mutations induced hSERT internalization independently of the allosteric binding site mechanism (58). In our experience, unintended consequences from mutation are common, and it is possible that mutagenesis of these sites alters the conformational structure of hSERT to promote a diminished palmitoylation and functional state, similar to hSERT binding by escitalopram, and warrants further analysis.

Currently, autism and its associated disorders are diagnosed exclusively through behavioral observation and assessment. It is prudent to discern the pathophysiological nature behind neurological/psychiatric disorders like autism so we can improve diagnostic strategies and therapeutic development. The data presented here provides an explanation for the reported efficacy of escitalopram in autistic patients that help attune undesired symptoms like anxiety. Although most autistic individuals don’t have hSERT coding variants like F465L, it is possible that other idiopathic mechanisms function to enhance hSERT palmitoylation with subsequent consequences, that may be sensitive to downregulation with SSRIs in a similar manner. Consistent with most somatic disorders, it is likely that psychiatric disorders have a multifactorial basis. Although many individuals may be pre-disposed to development of these disorders through altered genetics or epigenetics, it seems likely that internal physiologic changes induced by the environment may be responsible for triggering their development. We view the contribution of these studies as a platform to deepen excitement and exploration into how dynamic post-translational modifications can be used to explain the myriad of outcomes associated with all idiopathic neurological and psychiatric disorders, that may not harbor a genetic explanation. Likewise, we envision a future where research is directed toward uncovering how PTMs alter MAT structural conformations that promote or discourage subsequent functional consequences, or vice versa, in the context of induction by physiological stimuli and modulation of MAT structure by therapeutics. Deepened understanding of these complex molecular processes will clarify differences observed in the pathophysiology of neurological diseases and provide a platform for designing therapeutics that can target changes in MAT PTMs to rectify their underlying pathogenesis.

## LIMITATIONS

Tests assessing L550V for surface expression, transport capacity, and reactivity to inhibition of palmitoylation was not assessed. In addition, further characterization is necessary for internalization rates, degradative mechanisms, the impact of cysteine-mediated substitutions, and other SSRIs on these processes.

## CONCLUSIONS

The molecular data presented here outlines the implications of a novel, dynamic post-translational mechanism recently discovered in hSERT regulation in the context of the clinically relevant autism spectrum disorder. Here, we reveal that two hSERT coding variants associated with ASD, F465L and L550V, demonstrate enhanced palmitoylation, and that this process in F465L hSERT is sensitive to inhibition with 2-bromopalmitate or escitalopram. Furthermore, inhibition of this process coincides with a decrease in previously enhanced surface expression and transport capacity to basal WT levels. Collectively, these data offer a molecular pathogenic-to-therapeutic continuum for serotonergic-mediated autism spectrum disorders where a dysregulated process (enhanced hSERT palmitoylation) promotes functional outcomes (enhanced hSERT surface expression and transport capacity), and that this process may be remediated by a commonly used therapeutic (inhibition of hSERT palmitoylation by escitalopram) and thus return of these disordered functional outcomes (and likely clinical manifestations) to basal WT levels.

## ABBREVIATIONS

ABE: Acyl-biotinyl exchange
ANOVA: Analysis of variance
AP: Alkaline phosphatase
ASD: Autism spectrum disorder
CNS: Central nervous system
DAT: Dopamine transporter
DHHC: Palmitoyl acyltransferase
DMEM: Dulbecco’s modification of Eagle medium
DMSO: Dimethylsulfoxide
DSM-4: Diagnostic and Statistical Manual of Mental Disorders, Fourth Edition
DTT: Dithiothreitol
EDTA: Ethylenediaminetetraacetic acid
ER: Endoplasmic reticulum
Esc: Escitalopram
G418: Geneticin
HBSS: Hank’s balanced salt solution
HEK-293: Human embryonic kidney-293
HEPES: 2-[4-(2-hydroxyethyl)piperazin-1-yl]ethanesulfonic acid
hSERT: Human serotonin transporter
KCl: Potassium chloride
KRH: Krebs-ringer HEPES
LSD: Least significant difference
MAT: Monoamine transporter
mRNA: Messenger ribonucleic acid
NaCl: Sodium chloride
NaF: Sodium fluoride
NEM: N-Ethylmaleimide
NET: Norepinephrine transporter
NH_2_OH: Hydoxylamine
PBS: Phosphate buffered saline
PKC: Protein kinase C
PKG: Protein kinase G
PMSF: Phenylmethylsulphonyl fluoride
PTM: Post-translational modification
PVDF: Polyvinylidene fluoride
p38 MAPK: p38 mitogen-activated protein kinase
RIPA: Radioimmunoprecipitation assay
SB: Sample Buffer
SDS-PAGE: Sodium dodecyl sulfate-polyacrylamide gel electrophoresis
SERT: Serotonin transporter
SLC6: Solute carrier 6 family of transporters
SSRI: Selective serotonin reuptake inhibitor
US: United States
WT: Wild type
2BP: 2-Bromopalmitate
5HT: Serotonin

## ACKNOWLEDGEMENTS

We would like to thank Dr L Keith Henry and Dr Danielle E Krout for thoughtful discussion and kindly providing the SERT autism coding variant expressing cell lines used in this study.

## AUTHOR CONTRIBUTIONS

**CRB:** conceptualization, methodology, investigation, data curation, visualization, formal analysis, writing – original draft, writing – review and editing

**JDF:** conceptualization, methodology, data curation, funding acquisition, visualization, supervision, project administration, writing – review and editing

## DATA AVAILABILITY

The datasets used and/or analyzed during the current study are available from the corresponding author(s) on reasonable request with statistical analysis results included with this published article’s supplementary information files.

## DECLARATIONS

### Ethics approval

No animals were used in preparation of the current article.

### Consent for publication

Not applicable.

